# BudFinder: A Masked Auto-Encoder Vision Transformer Framework for Yeast Budding Detection

**DOI:** 10.1101/2025.11.04.686463

**Authors:** Phuc Nguyen, Zahra Mousavi Karimi, Adrian Layer, Markus Wan, Hetian Su, Jeff Hasty, Nan Hao

## Abstract

Yeast replicative lifespan is a crucial part of aging research, yet its quantification remains labor-intensive and time-consuming, particularly when using time-lapse imaging and microfluidics. Manual counting methods for cell division events are prone to bias and inefficiency, while existing automated approaches often require extensive annotated datasets. These limitations hinder the adaptability of such tools across different microfluidic setups. To address these challenges, we propose a versatile image analysis approach that accurately detects yeast cell division events. To reduce the burden of requiring a large cell division annotated dataset, we pretrained a Masked Auto-Encoder on large-scale segmented yeast cell images. This substantially reduced the annotated data needed to train the transformer model for detecting cellular division events. Additionally, the model is trained directly on budding event detection, circumventing reliance on arbitrary heuristics, such as changes in cell area. By leveraging self-supervised pretraining, we reduced the training data requirement to fewer than 50 mother cells (∼1,000 divisions), representing a >5-fold reduction compared to prior methods while maintaining comparable accuracy.

**Author Summary:** Our work addresses a longstanding challenge in in live-cell time-lapse microscopy analysis, namely, automating cellular division tracking while minimizing the amount of training data required. Traditionally, scientists identify each division event by manually inspecting thousands of time-lapse images, a process that is both tedious and prone to bias. While automated tools exist, they often require large amounts of annotated data to work effectively, limiting their use across different experimental setups. To overcome these barriers, we developed BudFinder, which can recognize and track cell divisions with far less training data. Using yeast replicative aging data as an example, we first trained a model to understand what a yeast cell “looks like”, using tens of thousands of segmented yeast cell images entrapped in our custom-built microfluidic device. Then, we taught it to detect budding events directly from time-lapse movies. This approach reduces the need for manual labeling by more than five-fold compared to previous structures in place, while maintaining accuracy comparable to existing methods. By making high-throughput analysis of cellular division more accessible, our work paves the way for faster and more scalable quantification of cellular dynamics.

## Introduction

Recent advances in live-cell time-lapse microscopy have enabled researchers to continuously monitor the dynamic behaviors and molecular states of single cells over long timescales. Although these techniques reveal rich temporal phenotypes, analyzing such large image datasets remains a major challenge, as it requires segmentation, tracking, and quantification of individual cells across thousands of frames.

For example, the budding yeast *Saccharomyces cerevisiae* serves as a model for studying replicative aging in mitotically active cell types such as stem cells (1–3). The total number of daughter cells each mother cell produces before ceasing to divide is defined as its replicative lifespan (4–6). By following single mother cells over time, researchers can link molecular dynamics to lifespan outcomes.

Capturing these trajectories requires imaging individual cells every few minutes for several days to record each budding event and monitor fluorescent reporters of gene expression (7–11). This is achieved by integrating microfluidic platforms, which trap hundreds of mother cells in parallel side channels, with high-resolution time-lapse microscopy (12–17). This setup keeps each mother cell stably positioned in the imaging field for its entire lifespan while continuously flushing away daughter cells, resulting in detailed time-lapse movies that capture the full course of replicative aging (Figure 1A,B).

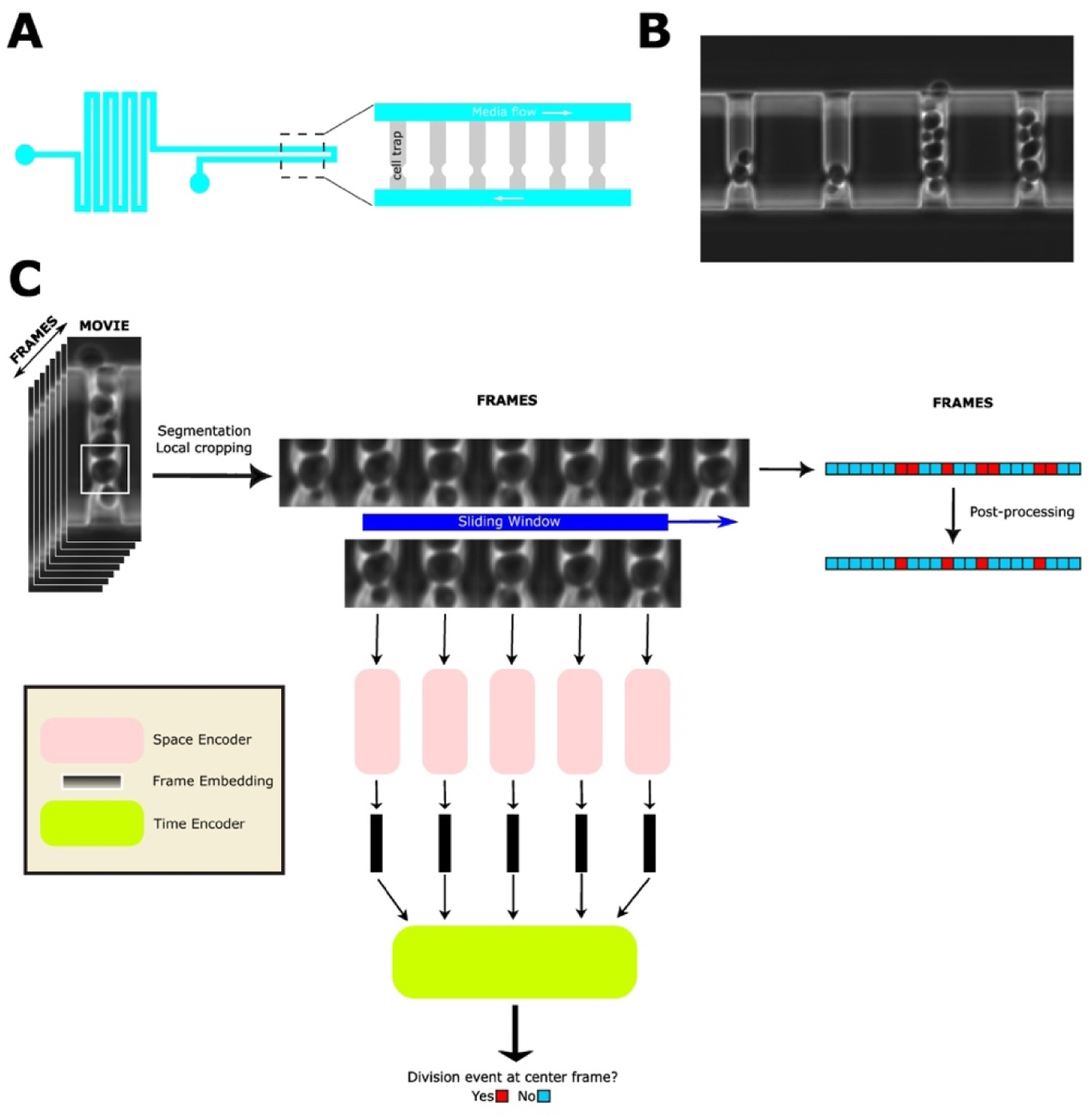

However, the data volume is enormous: a single 100-hour phase contrast run and a 15-minute sampling interval yields approximately 400 images per cell. Even with a lower limit of 100 mother-cell traps, a single experiment can yield thousands of raw images which must be managed before any preprocessing or quality-control steps can begin.

Traditionally, analysts comb through these stacks frame-by-frame, hand-annotating the exact moment of every bud emergence so that lifespan curves and cell-cycle metrics can be reconstructed (18). At roughly 20 to 30 buds per cell, this translates to tens of thousands of manual annotations per dataset, which is an effort that is both time-consuming and prone to subjective drift. In short, while time-lapse microscopy coupled with microfluidics unlocks unprecedented single-cell resolution, the manual curation that it demands throttles high-throughput investigation into aging mechanisms. Emerging deep-learning tools provide a promising avenue to break this bottleneck by automatically detecting cell division events directly from raw image sequences, and opening the door to much larger, statistically powerful inferences. Recent studies demonstrate that modern deep-learning architectures can match human annotators in identifying budding events while operating far more efficiently (19). For instance, YeastMate builds on Mask R-CNN, adding a custom head that simultaneously segments yeast cells and classifies lifecycle transitions (20). Similarly, DetecDiv leverages the mother-machine platform by coupling a U-Net encoder with a temporal attention module to classify each frame into six distinct states: unbudded, small budded, large budded, dead, empty trap, or clogged trap. This frame-by-frame classification is then used to reconstruct replicative lifespan trajectories (21). However, these approaches still face important limitations: they require large volumes of annotated data, which are costly and labor-intensive to generate, and involve complex multi-class classification schemes.

To address these challenges, we developed BudFinder, a Transformer-based framework optimized for efficient and accurate detection of cell division events in time-lapse microscopy data. Notably, BudFinder reduces the classification space to only two biologically meaningful states, budded vs. non-budded, which minimizes annotation complexity. Moreover, BudFinder achieves this performance with a dramatically smaller training dataset: fewer than 50 mother cells (∼1,000 annotated divisions) were sufficient for training. By comparison, YeastMate relies on more than 3,600 annotated budding events (plus an additional 2,380 mating events), and DetecDiv requires tens of thousands of annotated frames: 28,000 for frame-level classification and 150 full lifespans (150,000 frames) to achieve their reported performance. Overall, BudFinder reduces the training data requirement by roughly 3- to 6-fold compared with prior methods while maintaining state-of-the-art accuracy, underscoring its statistical efficiency and practical scalability. To achieve division detection with minimal annotated data, we hypothesized that the substantial demand for training data arises because the model must first learn to recognize and interpret the concept of a “cell” within an image before it can be effectively trained to detect complex cellular events such as division. As a result, a neural network can first be trained to recognize cellular imagery independently of the division-detection task. To implement this, a Masked Auto-Encoder (MAE) model is pre-trained on tens of thousands of segmented images of yeast cells in the mother machine. An MAE randomly masks a large fraction of image patches and trains a vision Transformer to reconstruct the missing content; thereby learning both local texture and global morphology features that are well suited to downstream tasks such as bud-event detection, even without task-specific labels. Importantly, high-quality segmentation masks are a prerequisite for MAE pre-training. By leveraging recent advances in cellular segmentation tools such as Cellpose (22), yeast segmentation can be effectively conducted with very limited training data: 50 manually-annotated frames (randomly generated from representative movies) were sufficient to bootstrap Cellpose on our pipeline (see *Methods*).

Second, a lightweight temporal Transformer head is attached and fine-tuned on less than 100 annotated clips. Self-attention allows the model to weigh every patch in every frame against each other, capturing long-range dependencies that CNNs miss. Transformers excel at “global context” while CNNs are restricted to local features, which translates into higher recall for atypical or slow-forming buds (23).

Overall, our experiments demonstrate that the proposed MAE and Transformer framework cuts annotation effort by roughly five-fold, substantially lowering the barrier to high-throughput yeast replicative lifespan studies.

## Results

### BudFinder framework accurately captures budding events and important aging-related metrics in time-lapse movies

To assess the accuracy of our automated cell division detection pipeline, we began with the timelapse images and the tracked coordinates of each cell (Figure 1C). We then compared the predicted replicative lifespan (RLS) against manually annotated ground truth data across wild-type yeast cells. The predicted RLS closely matched the ground truth values, with a high coefficient of determination (R² = 0.927), indicating strong concordance between model predictions and human annotations (Figure 2A). We further evaluated the performance of the division detection model using F1 score with a 1-frame tolerance window. The trained model significantly outperformed an untrained model baseline, achieving an F1 score of approximately 0.8 compared to <0.1 for the naïve approach (Figure 2B). In terms of RLS prediction error, the standard deviation from ground truth was under 3.5 divisions, demonstrating the model’s ability to detect divisions with high temporal precision despite the spatial and temporal complexities observed in the timelapse imaging dataset.

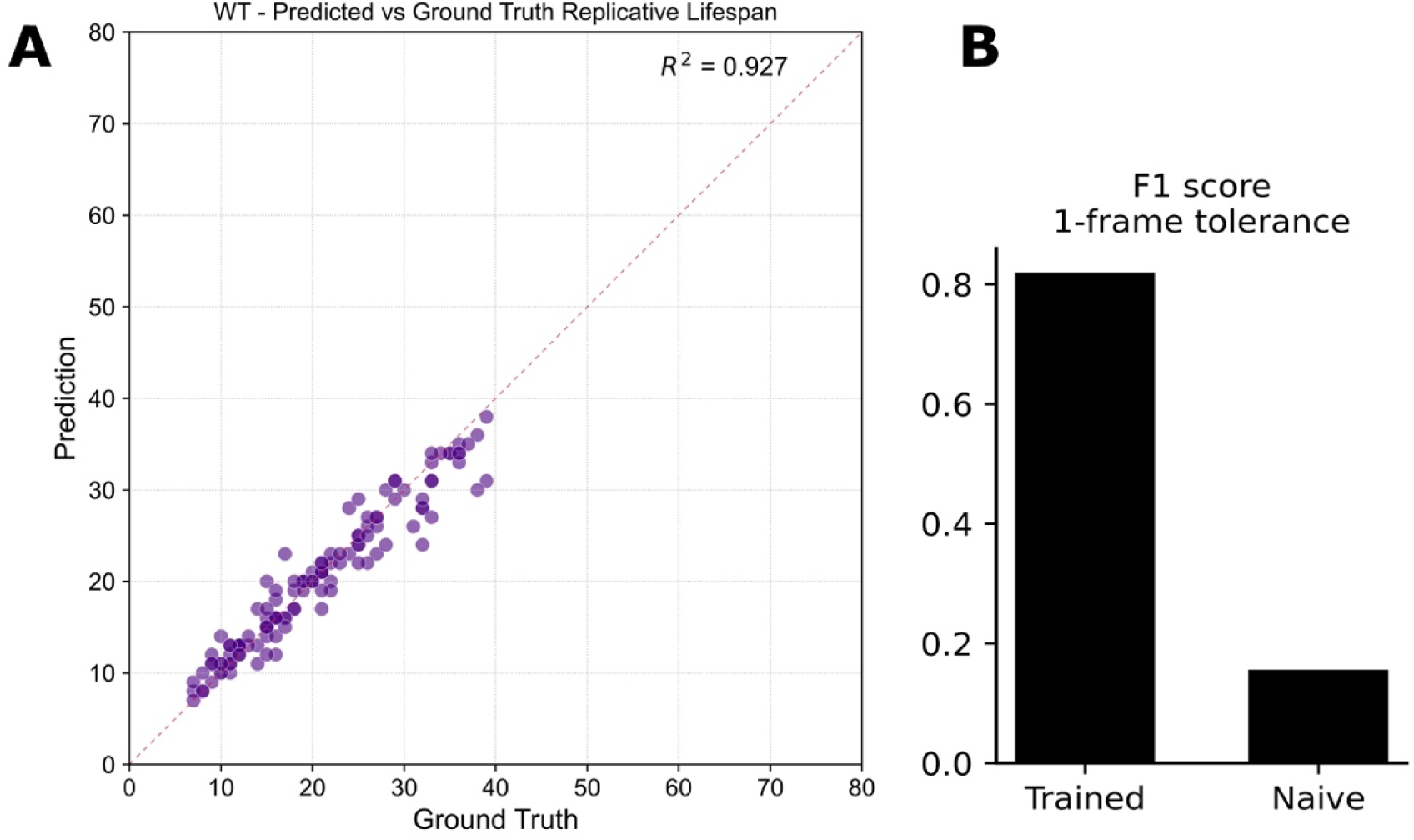

Given our framework’s ability to reliably detect budding events, we next asked whether its performance remained robust across the morphological differences characteristic of the two canonical yeast aging trajectories. In previous work, we stratified the wild-type population into two distinct phenotypes, referred to as Mode 1 and Mode 2 aging (28).

Mode 1 cells, roughly half of the cohort, switch from producing small round daughters to generating elongated daughters late in life and exhibit pronounced nucleolar enlargement and fragmentation, signaling nucleolar decline (Figure 3A, Top Left). Mode 2 cells, in contrast, continue budding small round daughters until death; their nucleoli remain morphologically unaltered, but they display progressive mitochondrial aggregation, a hallmark of mitochondrial dysfunction (Figure 3A, Top Right). Using these criteria, we labeled each cell track as Mode 1 or Mode 2 and evaluated the division-detection model accordingly. This partitioning allowed us to determine whether the model can recover replicative lifespan and cell-cycle dynamics despite the divergent morphological outcomes that define each aging mode. The predicted mean RLS values were consistent with the corresponding ground truth distributions across both aging modes, with no significant differences observed. This suggests that the model generalizes well across divergent aging trajectories and daughter cell morphologies (Figure 3B, 3C, Figure 1S, Figure 2S).

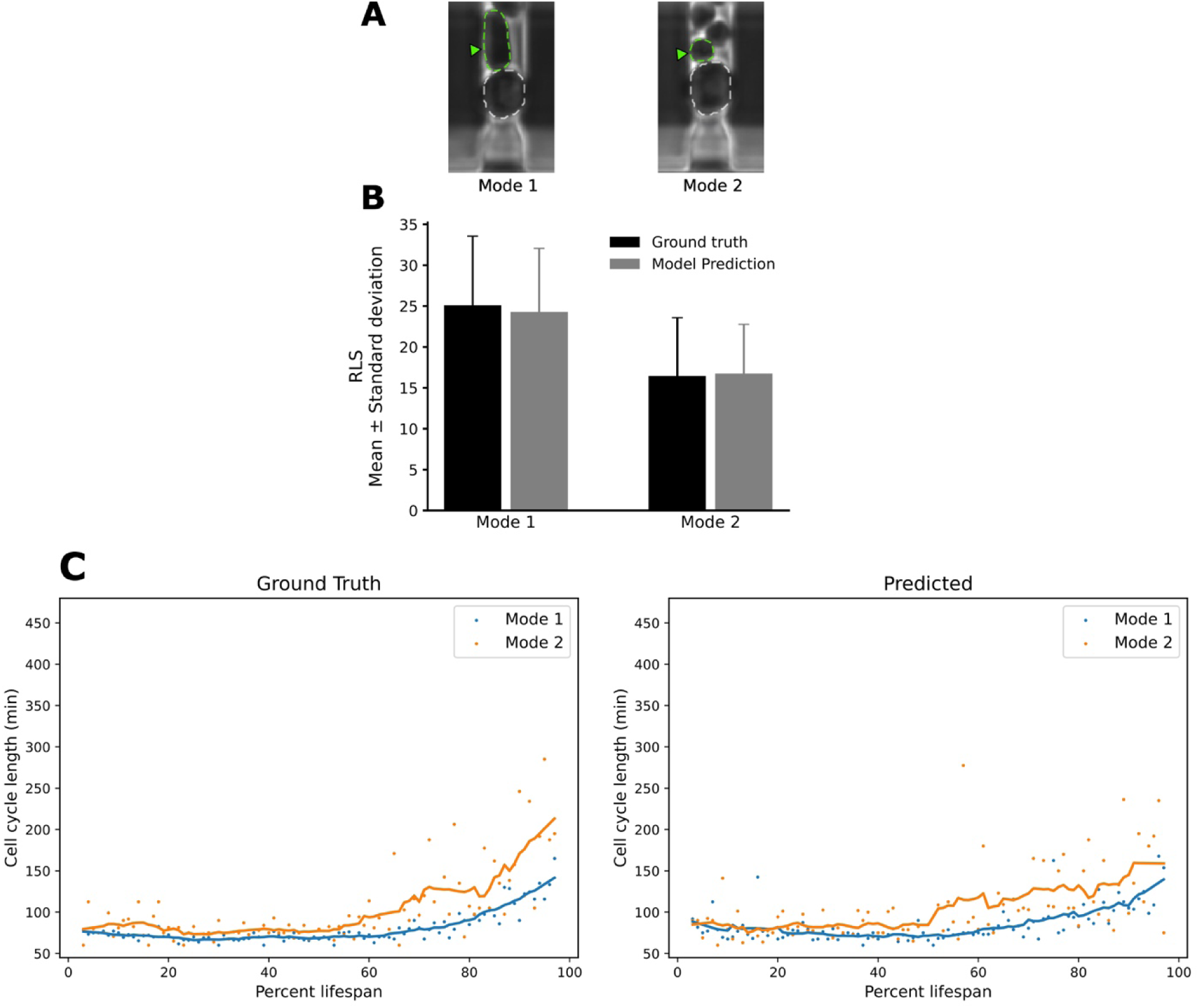

Overall, these results demonstrate that the division detection pipeline accurately reconstructs replicative lifespan and cell cycle dynamics, achieving high accuracy annotation performance while enabling high-throughput analysis of aging phenotypes.

### BudFinder generalizes across divergent aging phenotypes while maintaining accuracy at both frame and lifespan scales

We next tested whether the model maintains its accuracy in obtaining replicative aging information on short-lived or long-lived mutants in yeast: short-lived *sir2*Δ and a long-lived synthetic oscillator developed by Zhou et al. (24). We then assessed the performance between ground-truth data and the model at two complementary scales.

1. *Frame-level event detection:* For each strain, we compared the predicted and ground-truth budding timelines. Raster plots for 30 representative *sir2*Δ mother cells show that the model’s predicted division ticks (orange) align almost perfectly with the ground-truth ticks (blue) (Figure 4). Across 1,919 annotated frames, the model correctly identified 89.6 % of predictions occurring within ±2 frames of the manual labels, corresponding to an F1 score of 0.75 under a 1-frame tolerance (Table 1, Figure 5, Left). Similarly for the oscillator strain, the model captured 82.7% of 3,549 annotated budding events within ±1 frame, and 90.7% within ±2 frames of the manual calls, yielding an F1 score of 0.68 at the 1-frame tolerance (Table 1, Figure 5, Right). Event-timing precision therefore remains high despite both mutant’s distinct cell cycle dynamics.
2. *Replicative lifespan detection:* Summing the detected events yields a predicted lifespan for every mother cell. Scatter plots of predicted versus measured lifespans show tight agreement across both genotypes: R^2 = 0.882 for *sir2*Δ and 0.877 for the oscillator (Fig. 6). Notably, the regression slopes did not differ significantly from 1, demonstrating that the model generalizes without strain-specific recalibration. Together, these results confirm that our self-supervised/temporal-Transformer pipeline reproduces both the fine-grained cell-cycle timing and the coarse-grained lifespan statistics required for high-throughput aging studies.

**Figure 4.**
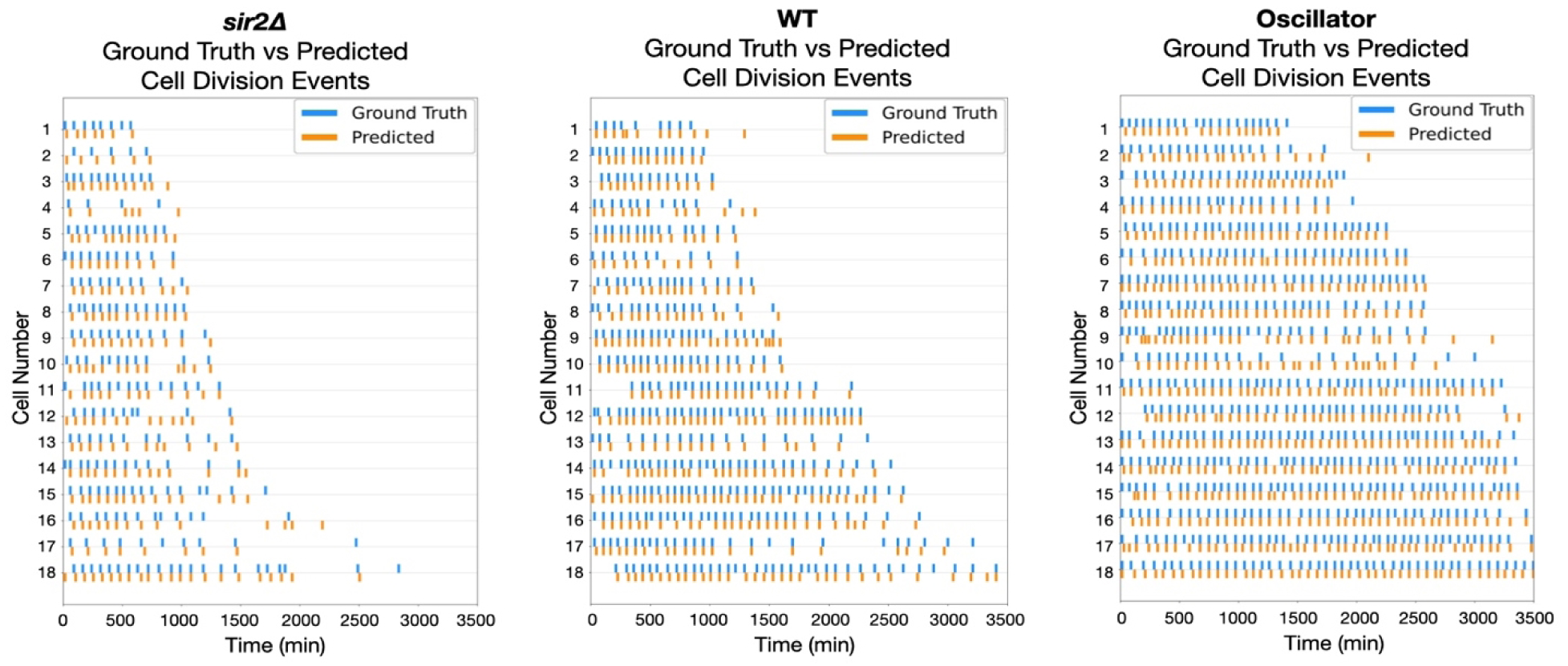
BudFinder reproduces manual budding timelines across diverse genetic backgrounds. Raster plots show ground-truth (blue) and model-predicted (orange) budding events for at least n = 15 representative mother cells per strain. The near-perfect superposition of orange and blue ticks across all three panels demonstrates that BudFinder preserves frame-level accuracy despite the distinct division frequencies and lifespan trajectories characteristic of each genotype, confirming its robustness to genetic perturbations.

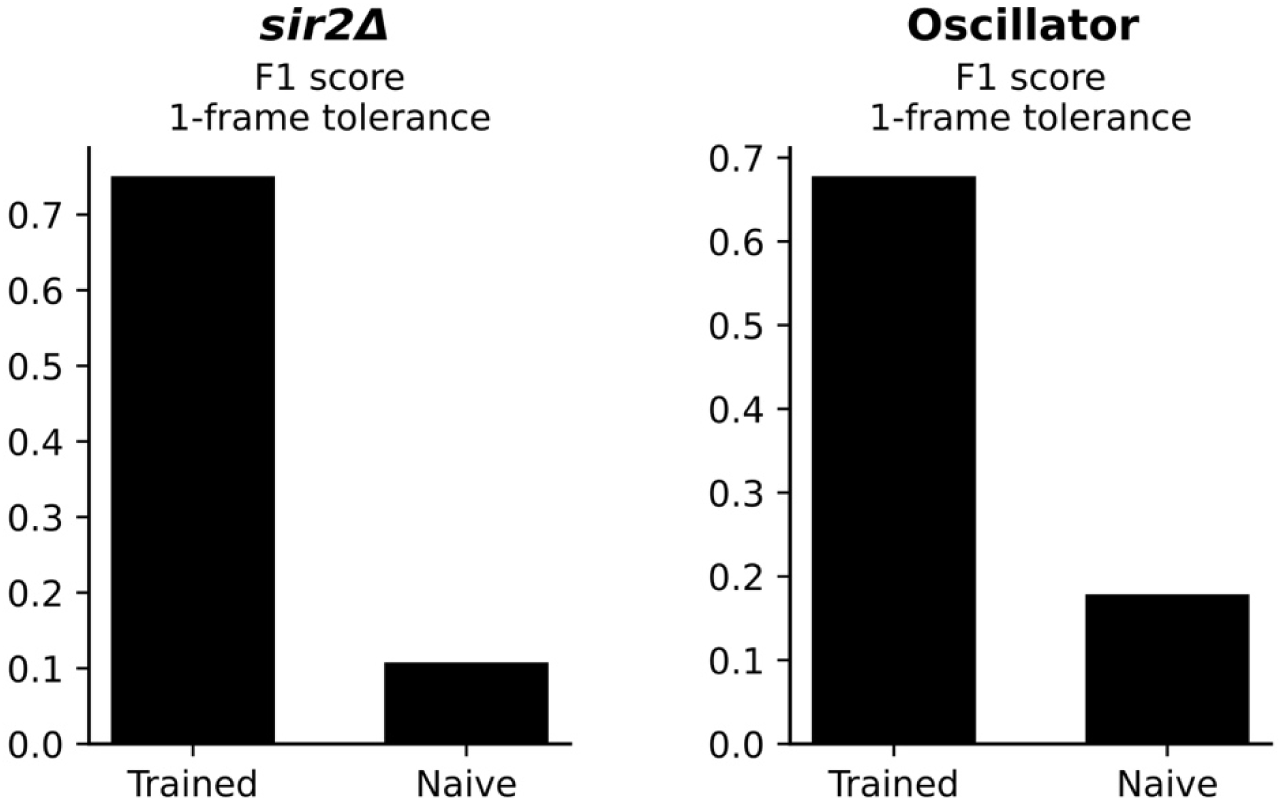

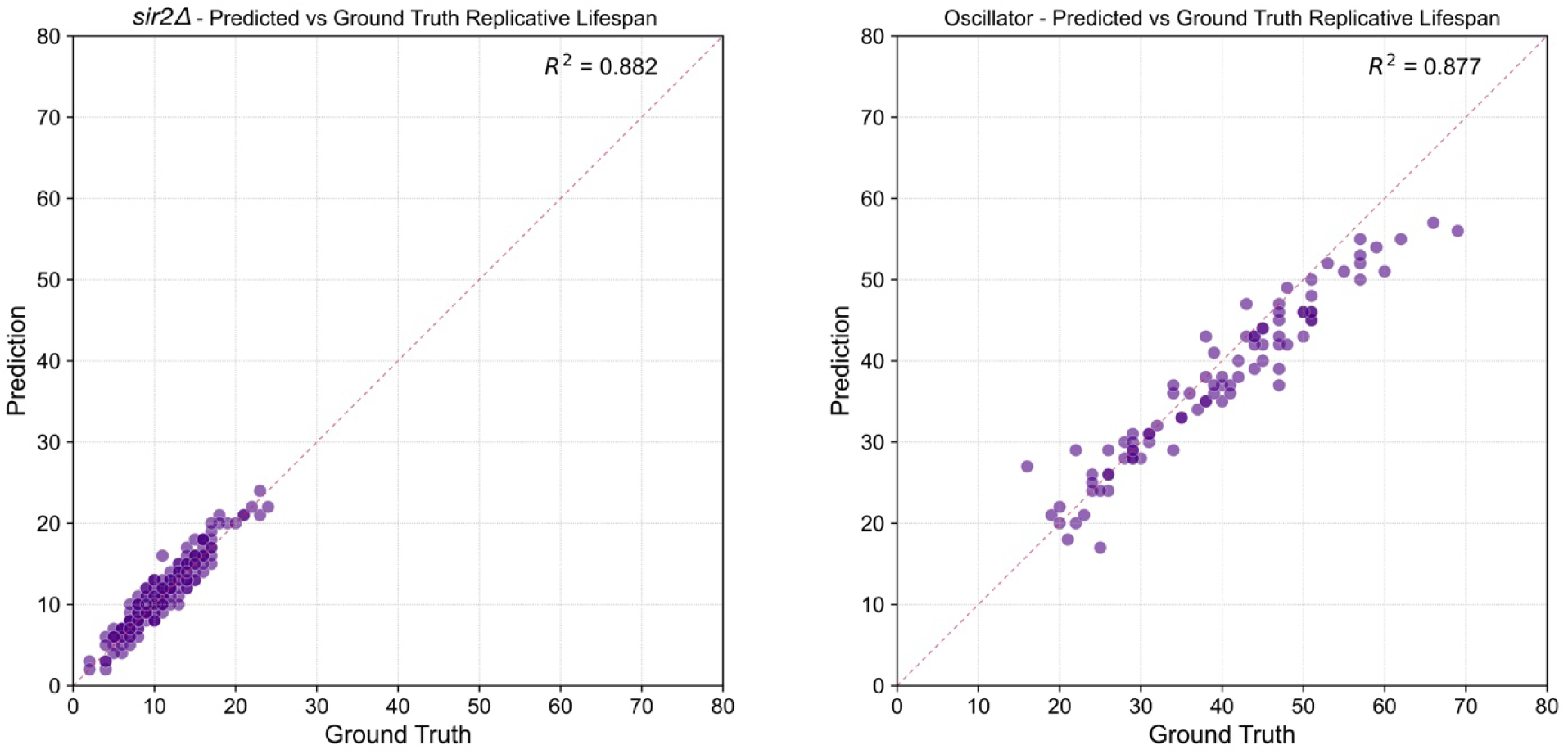

| Condition | Exact match | $\pm 1$ frame | $\pm 2$ frames |
| --- | --- | --- | --- |
| WT | 54.9% | 88.7% | 91.8% |
| Oscillator | 41.1% | 82.7% | 90.7% |
| *sir2Δ* | 32.5% | 78.4% | 89.6% |

Together, these results demonstrated the framework’s ability to effectively extract crucial aging-related information from complex timelapse movies by automated budding event detection. Once properly preconditioned, the MAE empowers the division-detection module to learn cell budding with reasonable training data and enables rapid, unmonitored examination, comparison, and analysis of aging experiments.

## Discussion

In this study, we present BudFinder, a Transformer-based framework that leverages MAE pretraining to enable accurate and efficient detection of yeast cell divisions. By combining self-supervised representation learning with a lightweight temporal Transformer, our approach reduces the annotated training data requirement by more than five-fold compared with existing methods, while maintaining state-of-the-art accuracy in reconstructing replicative lifespan trajectories. Importantly, BudFinder faithfully recapitulates budding timelines across wild-type and mutant strains. This demonstrates the model’s robustness to morphological variability, divergent aging modes, and altered cell cycle dynamics.

The novelty of our work lies in decoupling cell representation learning from division-event detection. We show that an MAE trained on raw yeast cell images provides morphology-aware embeddings that can be transferred to downstream classification tasks with minimal supervision. This strategy circumvents the need for vast, manually curated datasets, which is a persistent bottleneck in the field, and enables scalable, high-throughput analysis of time-lapse images.

While BudFinder demonstrates strong generalization, certain limitations remain. High-quality segmentation is essential for effective MAE pretraining, and pipeline performance may be reduced if segmentation masks are inconsistent across experimental setups. Moreover, although our framework reliably identifies division events, it does not explicitly monitor daughter morphology, limiting its ability to quantify features such as bud volume growth. Future work may integrate instance segmentation or multimodal architectures to extend BudFinder’s capacity beyond division counting.

The implications of BudFinder extend beyond yeast aging. Our framework highlights the utility of self-supervised pretraining in cellular image analysis, particularly for domains where annotated datasets are scarce. We envision that similar approaches could be applied to other single-cell imaging platforms, ranging from bacterial colony dynamics to mammalian stem cell differentiation. More broadly, the statistical efficiency of BudFinder opens the door to systematic, large-scale exploration of genetic and environmental factors that modulate replicative lifespan. By minimizing manual annotation and enabling large-scale, quantitative analysis, BudFinder provides a generalizable framework that advances live-cell imaging and deepens our understanding of dynamic cellular processes.

## Methods

### 1. Experimental Setup

#### Microfluidic Device Fabrication & Setup

The microfluidic devices and experiments were conducted according to previously established protocols (8,10,24). In summary, a microfluidic device was fabricated using a 4-inch silicon wafer patterned with SU-8 2000 series photoresists following standard photolithography techniques (Figure 1A). PDMS devices were then cast by pouring a Sylgard 184 mixture onto the wafer molds, degassing under vacuum, and curing for at least 1 hour. The PDMS devices were then cured at 60C overnight. The PDMS replicas were cut, punched, cleaned, and irreversibly bonded to glass slides using oxygen plasma. Detailed protocols for microfluidic device fabrication can be found in (O’Laughlin et al., 2023). Once the devices were prepared, to perform the experiments, yeast cells were cultured in SC medium with 2% glucose until reaching an OD600 of 0.8. The microfluidic devices were vacuum-treated for 20 minutes, coated with 0.075% Tween-20, and then placed on a 30°C incubator-equipped inverted microscope. Media ports were connected to syringes with synthetic dextrose media containing 0.04% Tween-20, positioned to enable gravity-driven flow. Yeast cells were loaded into the device, followed by media tubing reconnection. Flow rates were maintained at ∼2.5 mL/day with waste collected in tubes (Figure 1B).

#### Single-Cell Aging Time-Lapse Microscopy

Time-lapse microscopy was performed using a Nikon Ti-E inverted fluorescence microscope equipped with an EMCCD camera (Andor iXon X3 DU897) and a Spectra X LED light source. Imaging was conducted with a CFI Plan Apochromat Lambda DM 60× oil immersion objective (NA 1.40, WD 0.13 mm). Phase images were captured every 15 minutes over a duration of 90 hours, with an exposure setting of 50 ms.

### 2. Model Architecture

#### Overview

Our model was trained in two phases. In the first phase, an MAE was trained to reconstruct single-cell yeast images, enabling the encoder to learn morphology-aware representations. In the second phase, a classification head along with the pretrained encoder was trained to detect budding events from temporal image stacks. A final post-processing module takes in frame-level classification of a movie and outputs the following: (i) total replicative lifespan for each mother cell, (ii) the timing of division events, and (iii) frame-wise probabilities of division across the entire lifespan.

##### 2.1 Data Preprocessing

Raw microscopy data were acquired as 90-hour multichannel nd2 recordings and converted to plane-wise TIFFs. Frames were aligned to correct stage drift, and segmentation masks for individual cells were generated using Cellpose (22). Frame-resolved positional and morphological information was then extracted with ictrack (25) and assembled into a phenotype table (CSV format) that stored mother cell position, area, aspect ratio, and flat-field-corrected fluorescence intensities. For each 15-minute frame, one row per mother cell was appended, yielding an analysis-ready dataset in less than 24h. This preprocessing workflow not only produced high-quality morphometric and fluorescence measurements suitable for immediate statistical analysis but also streamlined downstream representation learning and event classification.

##### 2.2 Masked Auto-Encoder Pre-training

This encoder model learns rich representations of yeast images by reconstructing from random fragments.

For MAE pretraining, the preprocessing pipeline was truncated after the cropping step. To generate inputs for model training, centered 224 × 224 px crops were extracted for each mother cell at every frame and resized to 64 × 64. These crops were then flattened into patch sequences and augmented with positional embeddings for input into the model. Our implementation follows the architecture described in *Masked Autoencoders Are Scalable Vision Learners* (26). The model comprises an encoder, a decoder, and learnable mask tokens. During training, each input image was partitioned into patches, and a random subset of patches was retained and passed through the encoder to produce a latent representation. The decoder then reconstructed the full image from this latent representation together with the learnable mask tokens (Figure 7A). Positional embeddings were added to all tokens to preserve spatial structure, consistent with the design of ViTs (27).

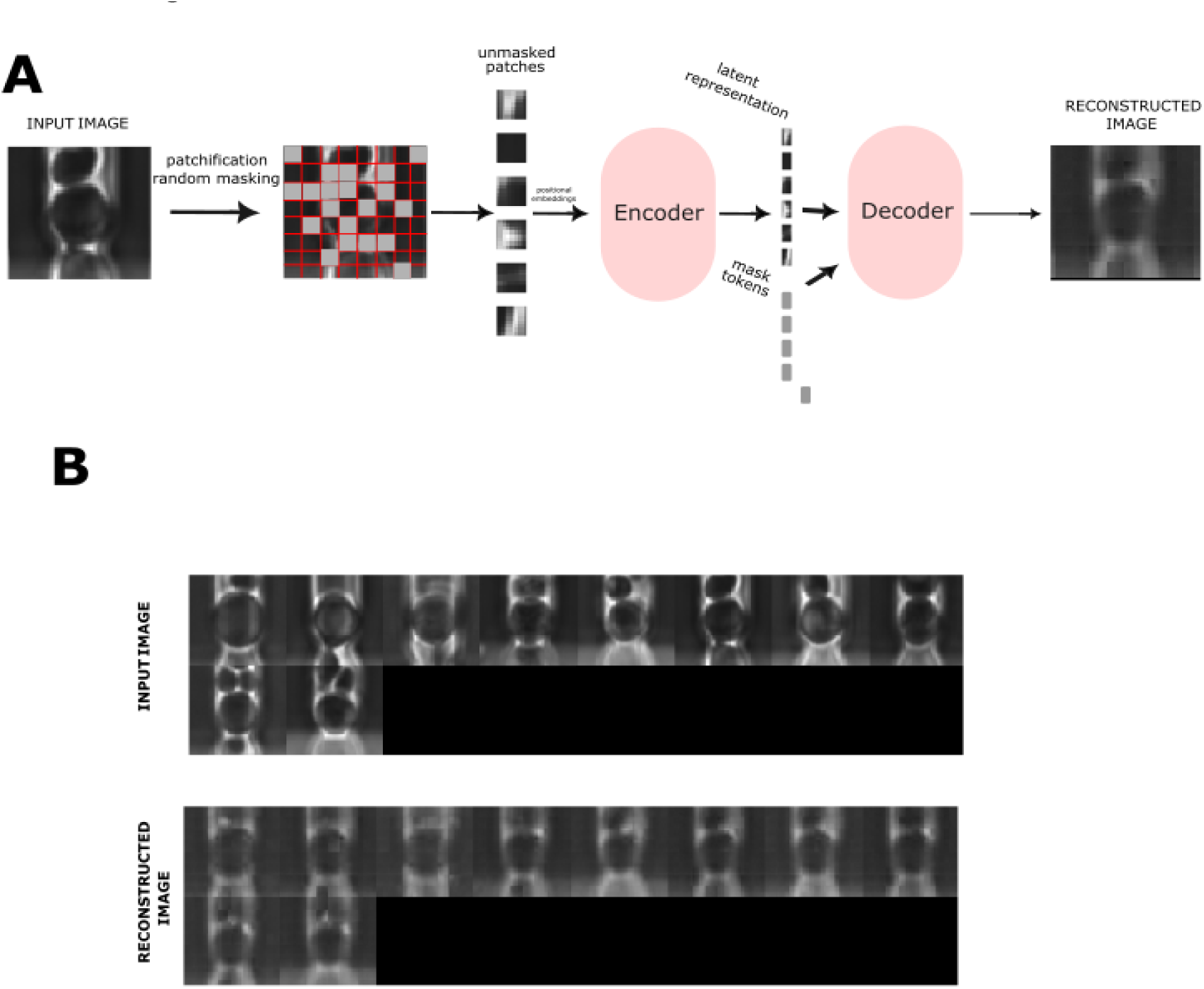

Training incorporated several strategies to improve stability and performance, including an exponential moving average (EMA) model for consistency, learning-rate schedules that combined warm-up with cosine annealing and periodic restarts, gradient accumulation to accommodate large batches, and the Adam optimizer with decoupled weight decay. The model was trained for over 500 epochs, and the best-performing checkpoint was retained for downstream applications (Figure 7B).

##### 2.3 Cell Division Detection Training

This model was trained to identify budding events from single-cell TIFF stacks. For division detection, temporal 224 x 224 px TIFF stacks were constructed by grouping 11 consecutive crops (the frame of interest plus five preceding and five subsequent frames). These stacks were obtained by running our preprocessing pipeline to completion. Each stack was patchified, linearly projected, and augmented with positional embeddings, analogous to the preprocessing used during pre-training. To capture temporal context, we further applied sinusoidal embeddings to encode the frame index within the sequence.

This model leverages weights imported from the encoder portion of the MAE, but is modified for classification tasks. The same vision Transformer encoder was retained, while the decoder was omitted. Unlike the self-supervised pre-training stage, no patches were masked so that the model could exploit all available information. Each frame was encoded independently, with a classification token prepended to generate a summary representation. These frame-level embeddings were then passed through an additional Transformer designed to integrate temporal information across the sequence. For prediction, the temporal Transformer aggregated features not only from individual frames but also across frame-to-frame transitions, enabling the model to weigh dynamic changes in morphology. Each frame produced a binary classification logit, which was converted into a probability via softmax. Predictions across a cell’s lifespan were aggregated, and post-processing was applied to collapse consecutive positive classifications into a single division event, thereby preventing over-counting (Figure 1B).

## Acknowledgement

This work was supported by NIH R01AG056440, R01AG086348, and R01AG093633.

## Data and materials availability

All data are available in the main text or the supplementary materials. The code from this work is available at https://github.com/haolab-ucsd/BudFinder/tree/main

## Supplemental Figures

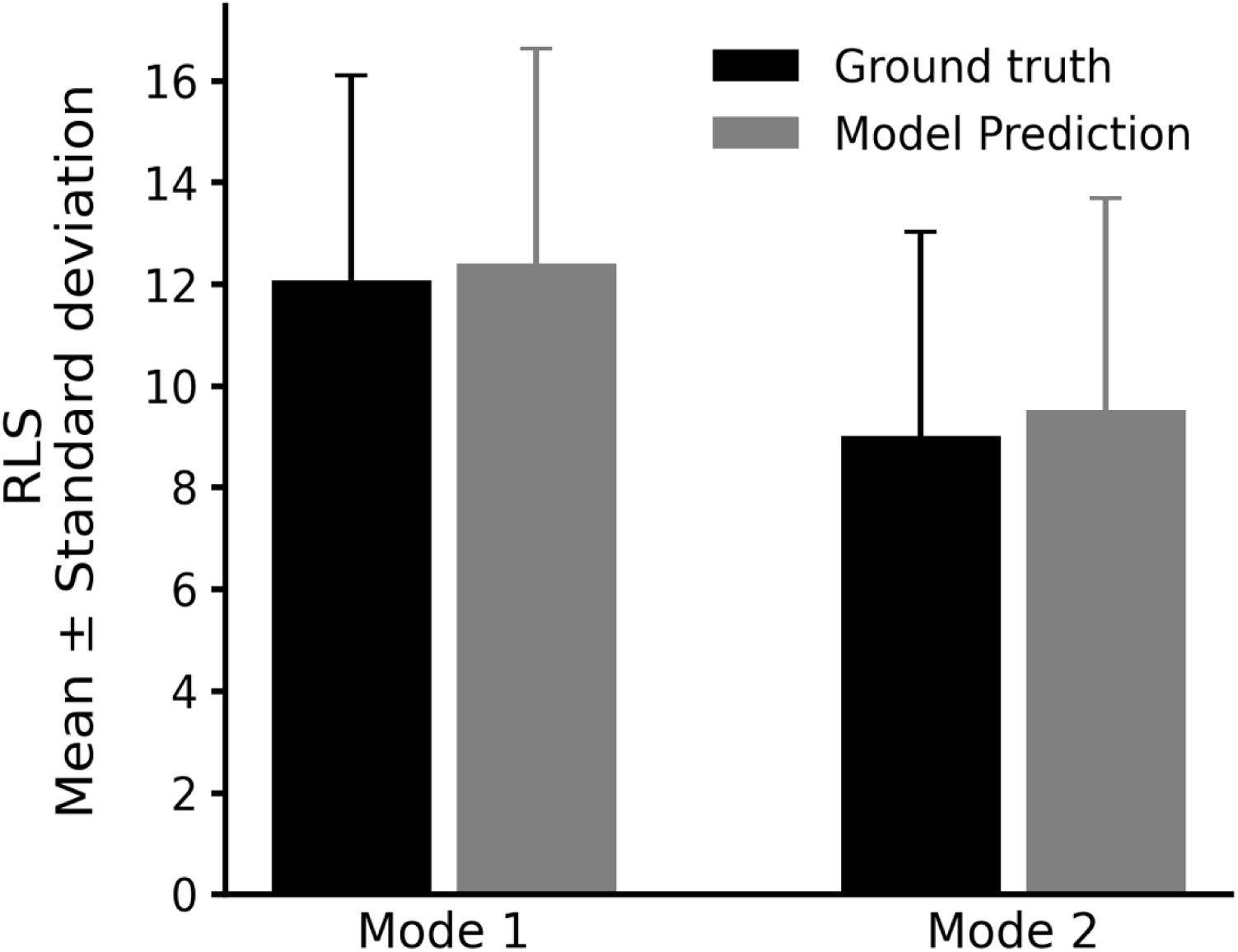

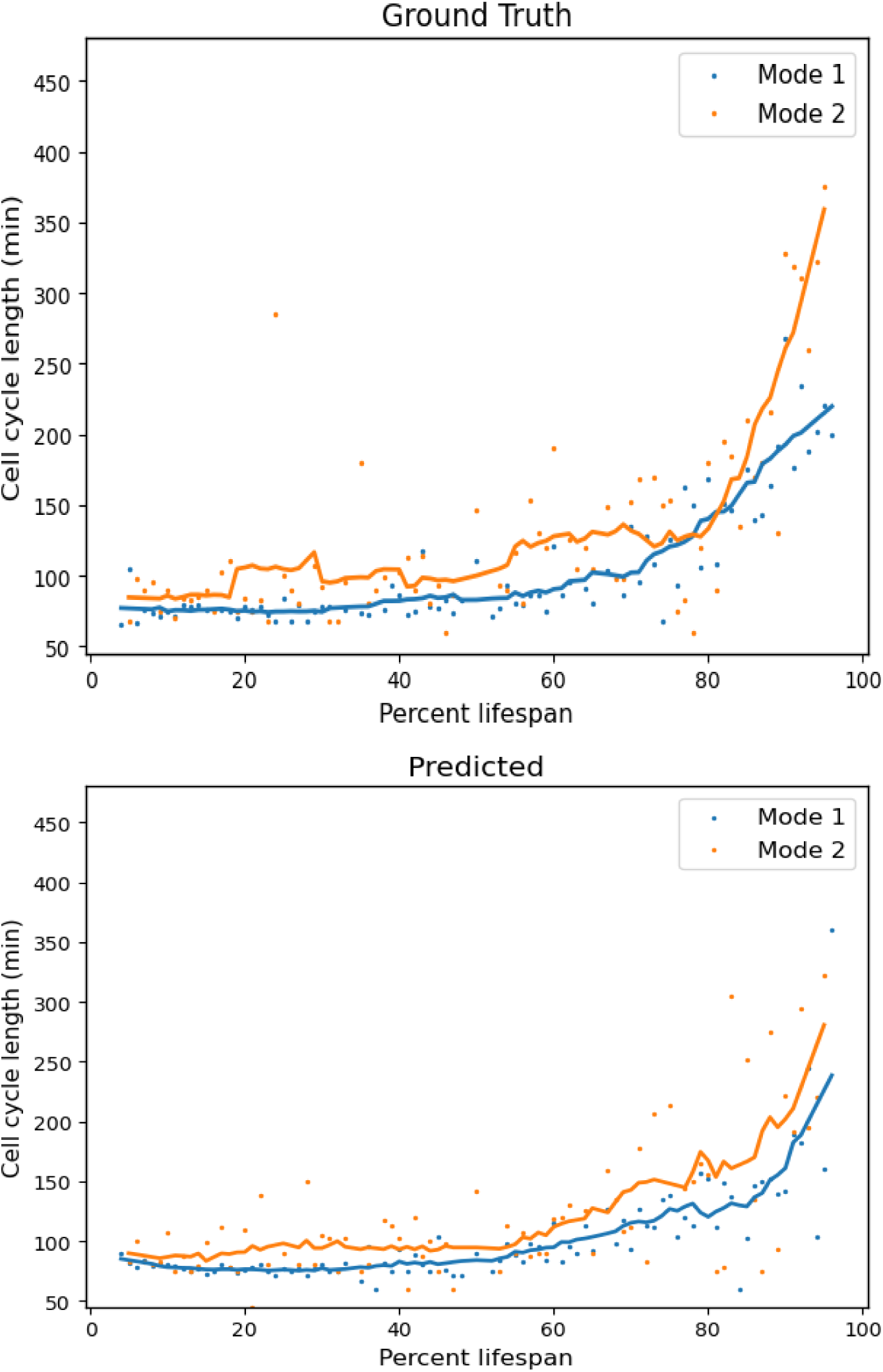

